# Aversion-induced drug taking and escape behavior involve similar nucleus accumbens core dopamine signaling signatures

**DOI:** 10.1101/2024.08.05.606651

**Authors:** Elaine M Grafelman, Bridgitte E Côté, Lisa Vlach, Ella Geise, G. Nino Padula, Daniel S Wheeler, Matthew Hearing, John Mantsch, Robert A Wheeler

**Affiliations:** Department of Biomedical Sciences, Marquette University, Milwaukee, WI 53233, USA; Department of Pharmacology and Toxicology, Medical College of Wisconsin, Milwaukee, WI, 53226

## Abstract

Dopamine release in the nucleus accumbens core (NAcC) has long been associated with the promotion of motivated behavior. However, inhibited dopamine signaling can increase behavior in certain settings, such as during drug self-administration. While aversive environmental stimuli can reduce dopamine, it is unclear whether such stimuli reliably engage this mechanism in different contexts. Here we compared the physiological and behavioral responses to the same aversive stimulus in different designs to determine if there is uniformity in the manner that aversive stimuli are encoded and promote behavior. NAcC dopamine was measured using fiber photometry in male and female rats during cocaine self-administration sessions in which an acutely aversive 90 dB white noise was intermittently presented. In a separate group of rats, aversion-induced changes in dopamine were measured in an escape design in which operant responses terminated aversive white noise. Aversive white noise significantly reduced NAcC dopamine and increased cocaine self-administration in both male and female rats. The same relationship was observed in the escape design, in which white noise reduced dopamine and promoted escape attempts. In both designs, the magnitude of the dopamine reduction predicted behavioral performance. While prior research demonstrated that pharmacologically reduced dopamine signaling can promote intake, this report demonstrates that this physiological mechanism is naturally engaged by aversive environmental stimuli and generalizable to non-drug contexts. These findings illustrate a common physiological signature in response to aversion that may promote both adaptive and maladaptive behavior.

## Introduction

Moment-to-moment changes in motivated behavior are required for an animal to successfully navigate an environment filled with often unpredictable adversity. Encounters with negative outcomes are marked by changes in an animal’s emotional state and decision making. This often promotes adaptive behavior to avoid, alter, or escape an unpleasant environment. Adversity is clinically relevant, as environmental stressors are frequently cited as a principle cause of relapse in those attempting to remain abstinent. Clinical data support these anecdotal observations^1–4^ and are consistent with a motivational theory that posits that aversive events promote drug seeking as a mechanism of coping^5,6^. Similarly, failure to effectively respond to stressors can result in the development or exacerbation of mood disorders^7^. Thus, understanding how aversive events are encoded in the brain and lead to changes in motivated behavior is essential for understanding and treating a variety of neuropsychiatric diseases.

A considerable amount of evidence indicates that dopamine (DA) signaling in the nucleus accumbens core (NAcC) is an essential component of the decision making process, where affective and associative information influence motivated behavior^8,9^. Both motivational state and DA signaling are modulated by aversive stimuli^10,11^. We and others have demonstrated that aversive stimuli and their predictors can decrease NAcC DA release^12–16^. Although motivated behavior is typically associated with increased activity in the dopaminergic system, there are situations in which DA reductions spur goal-directed action. One situation of potential clinical relevance is psychostimulant intake, in which pharmacological inhibition of DA activity, particularly in the NAcC, increases drug-taking^17–21^. This result has long been interpreted as the engagement of compensatory drug seeking behavior in response to artificial suppression of the DA system. However, it is not clear that this mechanism could be engaged when DA is inhibited by an ethologically relevant stimulus.

To determine whether environmentally induced suppression of DA can drive drug taking, we measured real-time NAc DA during cocaine self-administration sessions in which an acutely aversive 90 dB white noise (WN)^13^ was intermittently presented. We chose to examine intense WN because aversive auditory stimuli are well known to support negatively reinforced escape behavior^22,23^ and because aversive WN reliably alters DA signaling in a manner that resembles other aversive stimuli^13,15^. Then, to examine the generality of aversion-induced changes in DA to motivated behavior, we examined DA release patterns in an escape design in which rats could press a lever to terminate aversive WN.

## Methods and Materials

Detailed methodology can be found in **Supplemental Materials.**

### Subjects

Experiments were conducted on adult female and male Sprague-Dawley and Long-Evans rats (200-350g; Envigo) in an AAALAC-accredited vivarium following IACUC approved procedures. Procedures were conducted in operant conditioning chambers enclosed in sound attenuating cubicles (Med Associates). Group sizes for all analyses are described in Table S1.

### Surgical procedures

For the cocaine self-administration experiment, intrajugular catheters were implanted as previously described^24^. For photometry recordings, AAV5-hSyn-dLight 1.2 or 1.3b was injected into the NAcC and an optic fiber was implanted at the same site.

### Cocaine-only self-administration

After intravenous catheter surgery, recovery, and food restriction (90% body weight), 8 female and 7 male rats were trained in 6-7 daily 1-hr sessions to press one of two levers for a sucrose pellet on an FR1 schedule. Rats were then trained to self-administer cocaine on an FR1 schedule. The beginning of each cocaine self-administration session was signaled by the entry of both levers into the box and the illumination of two cue lights. Responses on the active lever resulted in a 2.6-s cocaine infusion (0.5 mg/kg/0.1 ml) accompanied by cue-light offset, active lever retraction, and a 20-s timeout period. Responses on the inactive lever resulted in no programmed consequences.

### WN testing during cocaine-only self-administration

After 12-14 daily 2-hr cocaine self-administration sessions, rats were tested with mild (55 dB) and intense (90 dB) WN. Each WN test session began with a 25-min self-administration period with no auditory stimulation. This was followed by 3 WN test cycles consisting of two 5-min WN presentations (WN “on” periods) separated by three 5-min quiet periods (WN “off” periods), for a total of 25 min (Fig 1A). Two intense test days and 2 mild test days (order counterbalanced) were conducted for a total of 12 WN exposures of each intensity. WN Test days were separated by 5 quiet test days with no WN presentations. The effect of WN on cocaine self-administration was assessed by subtracting the average number of infusions during the “off” periods from the infusions during the “on” periods. On quiet days, time-matched “on” and “off” periods were used for comparison. Due to technical issues, one male rat only received 1 intense test day, and another only received 4 quiet test days.

**Figure 1.**
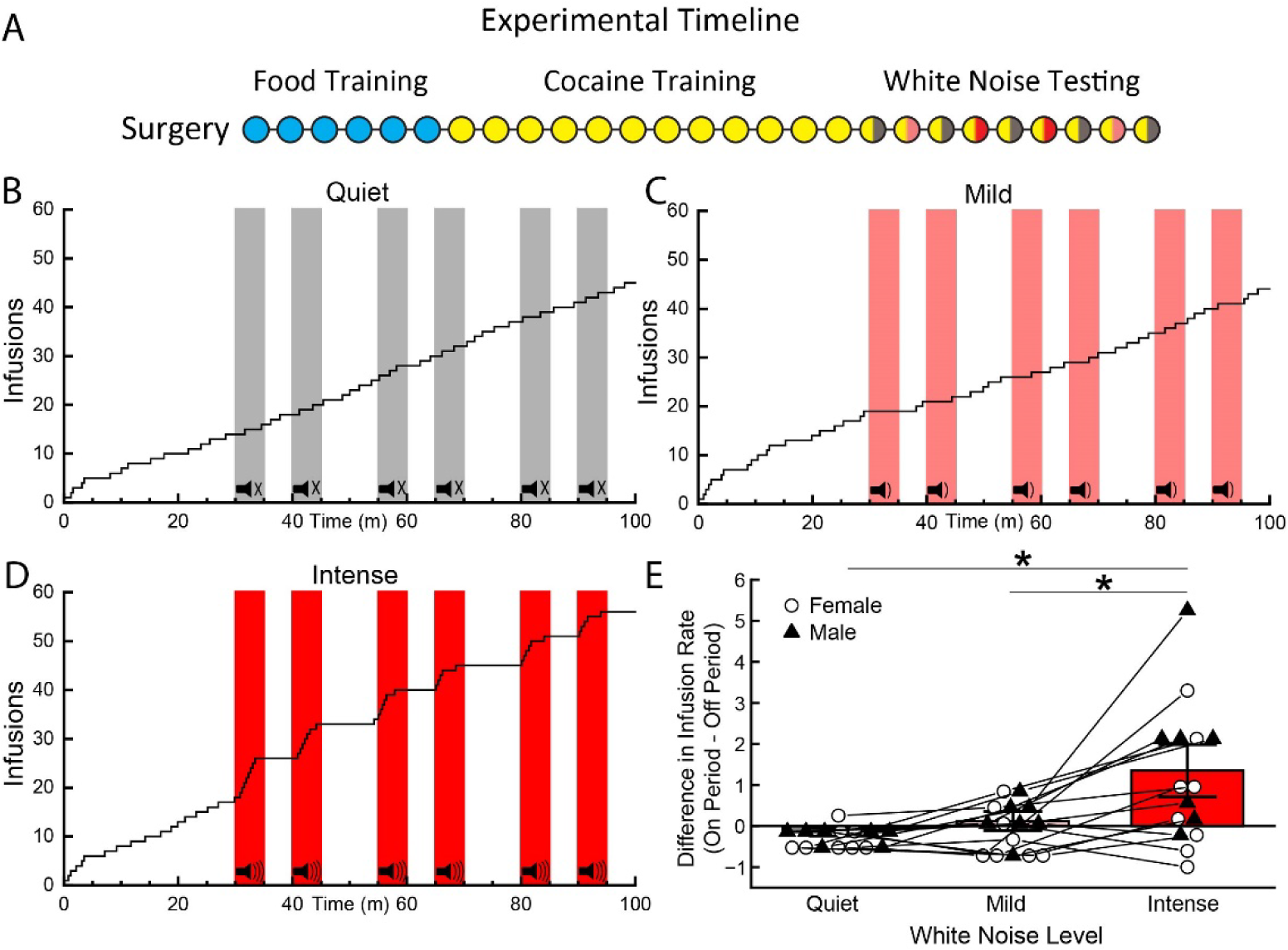
Presentation of Intense WN Promoted Cocaine Intake. (A) Experimental timeline. Following surgery, rats were trained in daily session for sucrose, then cocaine before test days in which different intensities of WN were presented during self-administration sessions. (B-D) Cumulative histograms depict operant responses for cocaine within single training sessions. Shading indicates noise presentation of 55 dB, 90 dB, or time-matched quiet period. (E) Mean difference in self-administration during noise on periods vs off periods. The presentation of intense WN (90 dB) caused an increase in cocaine intake compared to a comparable period, but this was not the case for the quiet test days and mild (55 dB) noise test days, F (2,13) = 10.27, p<0.01. Mean +/- SEM.

### Concurrent food and cocaine self-administration

A separate cohort of rats (7 male and 5 female) were trained to self-administer cocaine as described above with the following exceptions. Animals were initially trained to press two levers for sucrose pellets in daily sessions. Then all rats received 5 cocaine (0.8 mg/kg/0.2 ml infusion) self-administration sessions in which only one lever was present. Responses on this lever delivered only cocaine infusions for the remainder of the experiment. For the next 9-10 sessions, the second lever was reintroduced, and rats received simultaneous access to cocaine and sucrose pellets. Both reinforcers were presented on an FR1 schedule with the same response consequences and timeout. The responses were never mutually exclusive. Subjects then received 2 WN test sessions and 2 quiet test sessions. Each WN test day consisted of six 10-min intense WN on periods interspersed with 10-min WN off periods. Unlike previous tests, the test sessions began with a WN on period.

### Photometric recording during cocaine-only self-administration

A separate cohort of rats (6 male and 5 female) received IV catheter and dLight photometry surgeries and were trained to self-administer cocaine without concurrent food, as described above. Photometry recordings occurred on 4 nonsequential days featuring 2 quiet test days and 2 intense WN test days (counterbalanced). The test day procedure was identical to the cocaine-only test procedure described above. DA changes in response to WN were assessed by comparing the average Z-scored ΔF/F during the 60-s after noise onset (or matched “on” period during quiet days) to a baseline measure taken in the 60-s period immediately before noise onset. DA changes during cocaine infusions were assessed by comparing average Z-scored ΔF/F during the 5-s period immediately after the lever press to a baseline measure taken in the 2-s period starting 3-s before the lever press. To examine the relationship between DA and behavior, a difference score was calculated for each trial by subtracting the average DA during the 60-s baseline period from the average DA during the 60-s post onset period. Difference scores were averaged into 6 two-trial blocks and compared to the corresponding lever-press difference scores.

### Positive reinforcement

To examine DA signaling during positive and negative reinforcement, 23 female and 17 male rats received dLight photometry surgery. After recovery and food restriction, rats were trained in daily sessions to respond on two independently presented levers for a sucrose pellet on an FR1 schedule. Following 4 sessions, both levers were extended simultaneously and rewarded on independent VI30-s schedules. After 3 days, rats progressed to a VI90-s schedule for 6 sessions. Photometry recordings were taken on early and late day in VI90 reinforcement. Changes in DA were assessed by subtracting the average fluorescence in the 2- s period immediately following the lever press from the 2-s period immediately preceding the lever press.

### Negative reinforcement

A subset of rats (19 female, 16 male) that were recorded during PR also received NR testing. NR training sessions consisted of 60 daily trials in which responding on the left or right lever (counterbalanced) was reinforced by the termination of an aversive WN, while responding on the other inactive lever had no consequence. Rats were either trained with an intense or a mild WN. Each trial began with WN onset. After 5-s, both levers extended but responses on either lever had no consequence. Beginning 10-s after the onset of the WN, a response on the active lever resulted in the termination of the WN for 20-s. Responding on the inactive lever resulted in retraction of both levers for 5-s, but the WN persisted. If the rat failed to terminate the WN, it would persist for a total of 60-s, followed by a 6-s time-out period before the start of the next trial. After 5 days of training, 10 pseudorandomly interspersed 60-s quiet trials were introduced to the NR session for 3 training sessions. Photometry recordings occurred during both types of sessions. Animals were not included in the analyses if they did not yield data on one or more days of testing (Table S1). Changes in DA during responding were assessed by subtracting the average fluorescence in the 5-s period immediately following the lever press from the 2-s period immediately preceding the lever press. Changes in DA in response to WN were assessed by subtracting the average fluorescence in the 5-s period immediately following the WN onset from the 2-s period immediately preceding WN onset. Response probability was determined by dividing the number of responses by the number of presentations of the stimulus. Latency was defined as the average time to terminate WN.

### Analysis

Photometry data were extracted using Matlab, downsampled, and normalized as described in **Detailed Methods in Supplemental Materials**. Group and sex differences in cocaine self-administration, negative reinforcement, and DA were examined using ANOVAs and planned linear contrasts (PLC) using commercially available software (Statistica and R). The within-subjects relationship between DA and drug taking was characterized using repeated-measures correlation^25^. The between-subjects relationship between DA and negative reinforcement was characterized using Pearson correlation (Statistica).

## Results

### Intense WN Promoted Cocaine Intake

After cocaine self-administration training, rats were presented with different intensities of WN during daily test sessions to determine if these stimuli had an influence on drug intake patterns (Fig 1A-D). WN-induced changes in drug intake were summarized by difference scores, with a positive score reflecting elevated cocaine intake during WN relative to a pre-stimulus baseline. The presentation of intense WN (90 dB) had a stark effect on cocaine intake rates relative the quiet (no stimulus) and mild (55 dB) noise conditions (Fig 1E; ANOVA: F(2,13) = 10.27, p<.001). Cocaine intake was elevated during intense noise relative to mild noise (PLC: F(1,13) = 9.60, p=.009), as well as comparable time periods during the quiet test session (PLC: F(1,13) = 11.68, p=.005). WN did not affect responding on the inactive lever (Fig S2A; ANOVA: Fs<1.00, ps>.51).

### Intense WN Preferentially Promoted Cocaine Intake

During simultaneous sucrose and cocaine access, concurrent intake of food and drug was primarily limited to the initial portion of the session. After this, female rats began responding for food in a non-goal directed manner (responding compulsively without consuming the pellets earned), whereas males tended to stop responding for food entirely (Fig S2B; ANOVA: F(11,110) = 4.02, p<.001). No similar interaction was observed in cocaine taking (Fig S2C; ANOVA: Fs < 0.75, ps>.67). As such, we limited the WN test analysis to the first 20-min of the session. As with the previous experiment, we observed enhanced cocaine intake during intense WN (Fig 2A; ANOVA: F(1,10) = 20.33, p=.001). In contrast, food responding was not enhanced (ANOVA: F(1,10) = 3.17, p=.11), and was even suppressed in male relative to female rats (ANOVA: F(1,10) = 7.75, p=.02).

**Figure 2.**
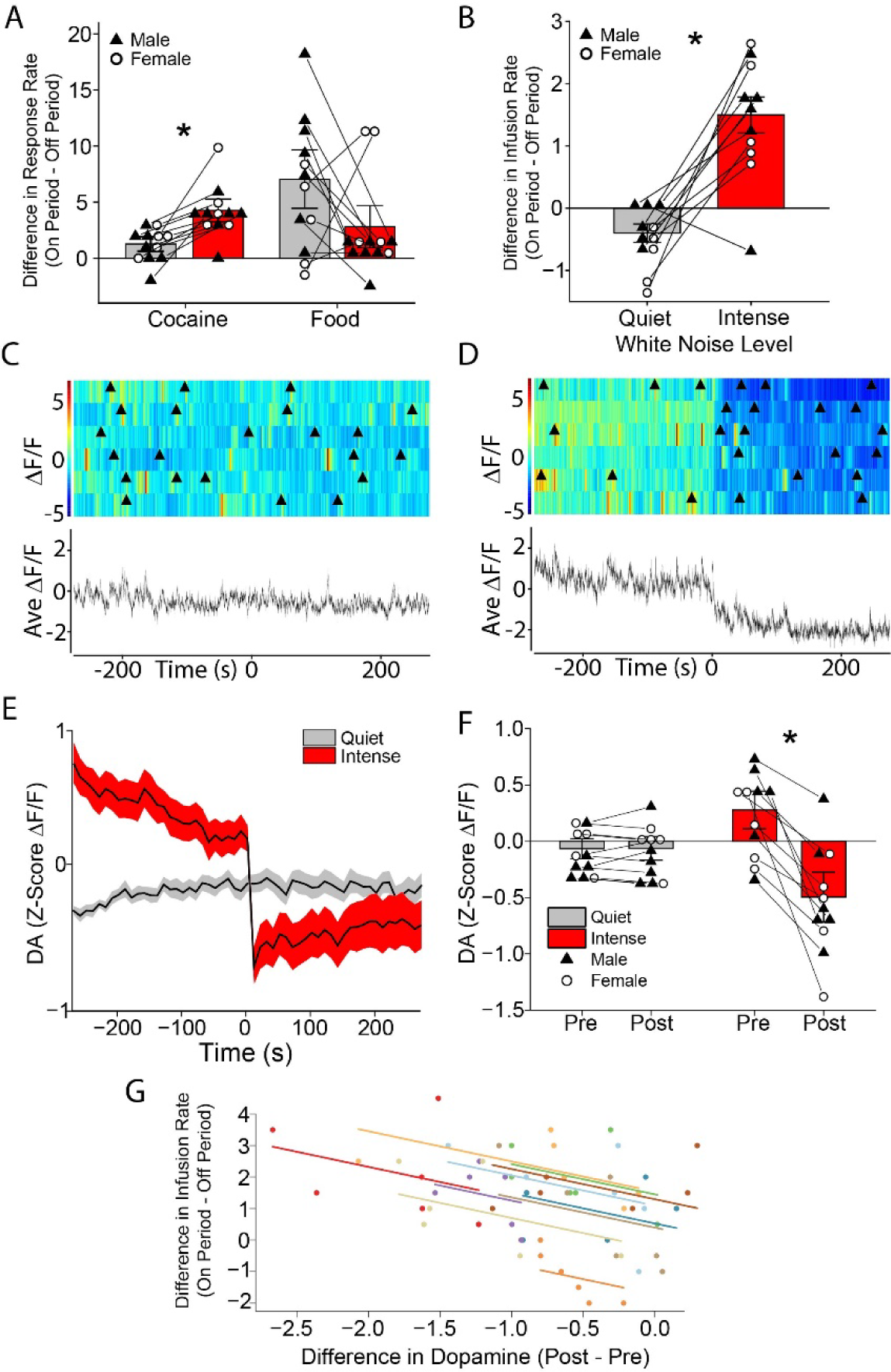
Intense WN Reduced Dopamine and Promoted Drug-Taking Behavior. (A) Mean difference in lever pressing during noise on periods vs off periods during concurrent food and cocaine access. The presentation of intense WN (90 dB) caused an increase in cocaine intake relative to quiet test days, F (1, 10) = 20.33, p < .01, but did not increase responding for food. Mean +/- SEM. (B) Mean difference in self-administration during noise on periods vs off periods during photometric DA recordings. The presentation of intense WN (90 dB) caused an increase in cocaine intake relative to quiet test days, F (2, 9) = 25.87, p<0.01. Mean +/- SEM. (C-D) Peri-event rasters depicting trial-by-trial changes in DA aligned to noise presentation or a comparable quiet time period. Triangles depict operant responses for cocaine. (E) Line graph depicting the mean time-averaged concentration change in DA in all animals aligned to the onset of Intense WN or a comparable quiet period. (F) Mean difference in DA concentration before (Pre) vs after (Post) onset of noise (red) compared to a comparable period on a quiet test day (grey), F (1, 9) = 24.20, p < .01. Mean +/- SEM. (G) Average noise-induced change in cocaine intake and DA (dots) for each subject (different colors) across six 2-trial blocks. Cocaine intake was greater on trials in which DA was more suppressed by intense noise, r_rm_ (49) = -0.40, CI [-0.61, -0.14], p < .01. Parallel lines depict fitted linear regression for each subject.

### Intense WN Reduced NAcC Dopamine and Promoted Drug-Taking Behavior

When DA concentration changes were examined during self-administration sessions, it was clear that the noise-induced increase in drug intake was aligned with a noise-induced reduction in DA (Fig 2B-G). The presentation of intense WN again caused an increase in cocaine intake relative to quiet test days (Fig 2B; ANOVA: F(2,9) = 25.87, p<.001). Simultaneously, intense WN presentation impacted dopamine concentration (Fig 2E,F; ANOVA: F (1,9) = 24.20, p<.001). WN reduced DA relative to the pre-WN baseline (PLC: F (1,9) = 28.39, p<.001), whereas DA was stable on quiet test days (PLC: F (1,9) = 0.05, p=.82). Similar to the previous test, WN did not affect responding on the inactive lever (Fig S2D; ANOVA: Fs < 2.00, ps>.19).

### NAcC Dopamine Release Inversely Correlated with Drug Taking

To further characterize the relationship between DA and behavior, we calculated difference scores (Post – Pre) for average DA levels during WN testing across 2-trial blocks and compared these measurements to drug taking during the corresponding trials (Fig 2G). A repeated-measures correlation revealed a moderate negative relationship, showing that the enhancement of drug-taking was greater on trials in which DA was more suppressed by intense noise, r_rm_(49) = -0.40, CI(-0.61, -0.14), p=.004.

### WN Suppressed NAcC Dopamine Elevations Aligned with Cocaine Self Administration

Operant responding for cocaine was associated with a transient elevation of DA concentration. However, this elevation was diminished when the response occurred during the presentation of intense WN (Fig 3A-C; ANOVA: (F(1,9) = 6.65, p=.03). Although DA levels were comparable during the pre-response baseline (PLC: F(1,9) = 2.11, p=.18), post-response DA was higher on quiet test days (PLC: F(1,9) = 7.15, p=.03). The inhibiting effect of WN on infusion-induced DA did not vary based on sex (ANOVA: F(1,9) = 0.05, p=.83), but the DA elevation associated with the lever press was greater in male rats (ANOVA: F(1,9) = 6.13, p=.04).

**Figure 3.**
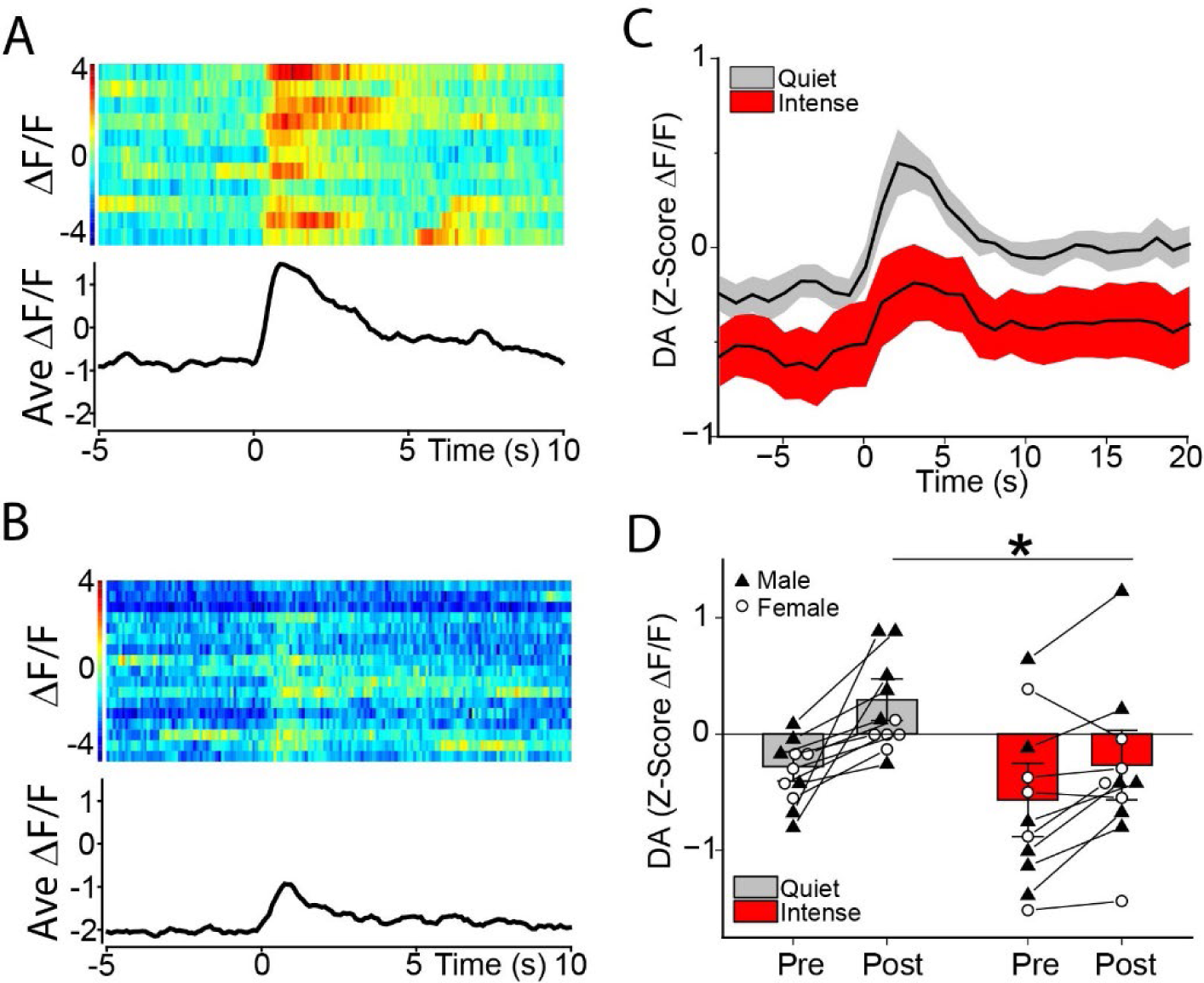
WN Suppressed Dopamine Elevations Resulting from Cocaine Self Administration. (A-B) Peri-event rasters depicting trial-by-trial changes in DA aligned to the operant response for cocaine during a quiet session in which noise was not presented (A) or during presentation of an intense WN (B). (C) Line graph depicting the mean time-averaged concentration change in DA in all animals aligned to the operant response for cocaine during a quiet session or an intense WN session. (D) Mean difference in DA concentration before (Pre) vs after (Post) operant response for cocaine in as session when noise was not presented (grey) vs during noise presentation (red), F (1,9) = 6.65, p<0.05.

### Intense WN Maintained Negatively Reinforced Behavior

In order to ensure robust lever pressing behavior to support negative reinforcement learning, rats were first trained to respond for food on a variable interval schedule (Fig 4A). After positive reinforcement training, food rewards were discontinued. Rats were presented with trials of either intense or mild WN and responses on one lever (active) terminated the noise while responses on the other lever (inactive) did not (Fig 4A). Across days of negative reinforcement training, rats that received intense noise maintained active lever responding at a higher rate than those that received mild WN (Fig 4B; ANOVA: F(8,248) = 8.27, p<0.001). On the last day of positive reinforcement training, responding on the active lever was similar in the mild and intense conditions (PLC: F(1,31)=0.49, p>0.49), but responding was lower in the mild condition on the final day of negative reinforcement (PLC: F(1,31)=31.14, p<0.001). A corresponding analysis of inactive lever responding revealed a downward trend in both conditions (ANOVA: F(8,248) = 200.77, p<0.001), and an interaction between sex, intensity, and training day (ANOVA: (F(8,248) = 2.11, p=0.035). In addition to responding more frequently on the active lever, rats in the intense condition also terminated the noise faster (Fig 4C; ANOVA: (F(1,37) = 58.33, p<0.001).

**Figure 4.**
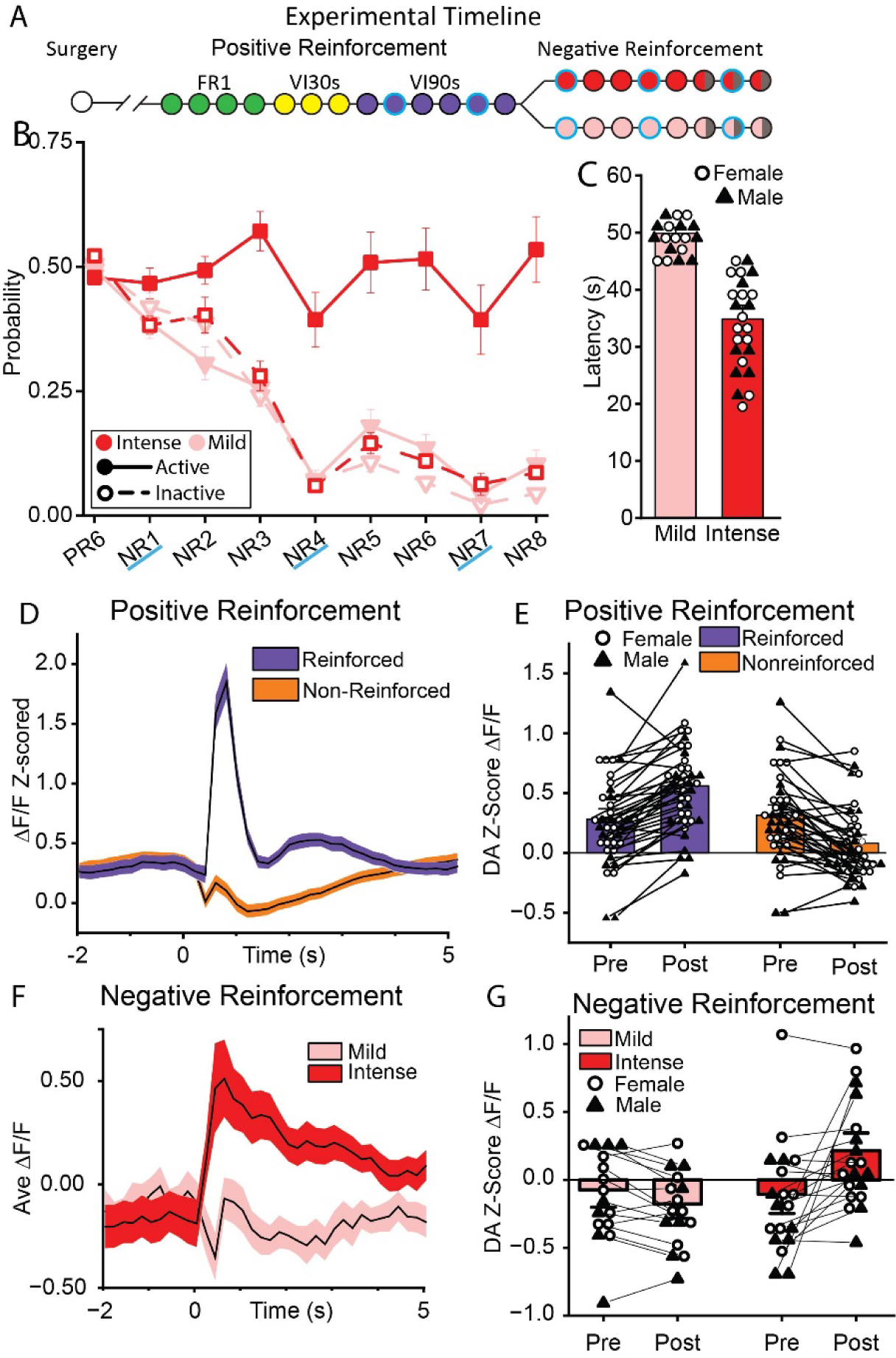
Intense, but not Mild, WN Maintained Negatively Reinforced Behavior. (A) Experimental timeline. Following surgery, rats were trained in daily sessions to respond for sucrose at various intervals, then transitioned to Negative Reinforcement wherein they received either mild (55 dB) or intense (90 dB) WN and were able to terminate it with a response on an active lever. (B) The probability of a response on the active (solid) or inactive (open) levers across days of training for rats receiving intense (red) or mild (pink) WN. On the last day of positive reinforcement, intense and mild animals made the same number of responses (F(1,31) = 0.49, p=0.489), but by the last day of negative reinforcement, mild-trained animals performed fewer responses than intense-trained animals (F(1,31) = 31.14, p<0.001). (C) The latency to WN termination under intense (red) and mild (pink) conditions. Rats receiving intense WN terminated the noise faster than those receiving mild WN (F(1,37) = 58.33, p<0.001). (D,E) Reward delivery elevated DA above pre-response concentrations (PLC: F(1,37)=52.77, p<0.001), whereas reward omission reduced DA (PLC: F(1,37)=31.18, p<0.001). (F,G) Mild and intense WN termination differentially regulated DA signaling, (ANOVA: F(1,30) = 17.32, p<0.001). Termination of intense WN increased DA, (PLC: F(1,30)=55.55, p<0.001). Termination of mild WN had no effect (PLC: F(1,30)=0.29, p=0.592).

### NAcC Dopamine Signaled Reward Delivery and Omission

Consistent with prior studies, reward and omission differentially impacted DA (Fig 4D,E). Reward delivery sharply elevated DA above pre-response concentrations (ANOVA: F(1,37)=52.77, p<0.001), whereas reward omission reduced DA (ANOVA F(1,37)=31.18, p<0.001).

### Intense WN Reduced NAcC DA During Negative Reinforcement Learning

DA signaling during WN presentations were recorded on the first day of negative reinforcement training to assess initial learning (Fig 5). Noise intensity modulated DA signaling (Fig 5D; ANOVA: F(1,30) = 20.16, p<0.01), with intense WN producing a clear reduction in DA (PLC: F(1,30)=26.24, p<0.001), and mild WN producing no reliable effect (PLC: F(1,30)=1.11, p=0.299). Although the behavioral task is quite different, WN modulation of DA was similar to that observed in cocaine self-administration. The suppression of DA by intense white noise was consistent across training days (Fig S3). WN termination, which served as the reward for negative reinforcement behavior, also clearly impacted DA signaling (Fig 4F,G; ANOVA: F(1,30) = 17.32, p<0.01). When rats successfully terminated the intense WN on the first day of NR, DA sharply increased (PLC: F(1,30) = 55.55, p<0.001). No similar effect was observed when rats terminated the mild WN (PLC: F(1,30)=0.29, p=0.592). Thus, the rewarding impact of WN termination in negative reinforcement had a similar neurochemical signature as food delivery in positive reinforcement.

**Figure 5.**
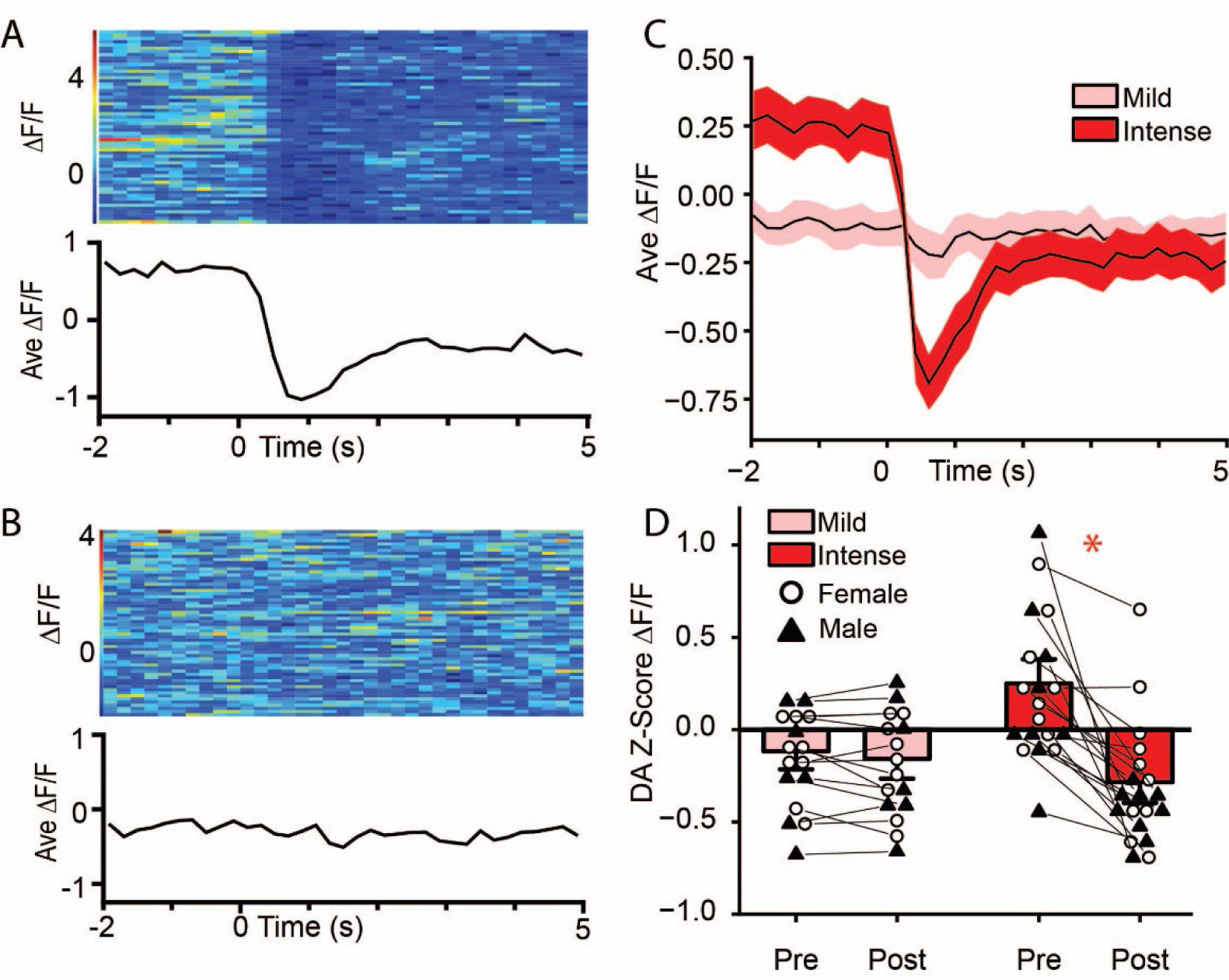
Onset of Intense, but not Mild, WN Reduced NAc Dopamine on the First Day of Exposure. (A-B) Peri-event rasters depicting trial-by-trial changes in DA aligned to Intense (A) or Mild (B) WN presentation. (C) Mean +/-SEM DA concentration timelocked to onset of Intense (red) or Mild (pink) WN on the first day of negative reinforcement. (D) Dopamine concentration in the 2s prior to (pre) vs 5s after (post) WN onset, (F(1,30)=26.24, p<0.001 for Intense; F(1,30) = 1.11, p=0.300 for Mild).

While mild WN did not inhibit DA or support negative reinforcement learning early in training, rats began to show a moderate but significant reduction in DA in response to the mild WN after a few days of exposure (Fig S3). This created an opportunity to determine if the decrease in NAc dopamine reflected a sensitization to the aversive properties of the mild WN such that the same stimulus could now support the acquisition of negatively reinforced behavior. Upon retraining, this mild stimulus that now reduced DA also supported negative reinforcement learning (Fig S4).

### Reductions in Dopamine Correlated with Enhanced Escape Behavior

Finally, we examined the relationship between DA signaling during initial learning of negative reinforcement and subsequent performance. Reductions in DA caused by WN (Post - Pre) on Day 1 were correlated with behavioral measures of escape on Day 8. That is, the magnitude of the initial DA reduction in response to WN was inversely correlated with escape rate on Day 8 (Fig 6A; Pearson’s *r*=-0.50*, p*=0.003). Initial DA responses were directly correlated with latency to escape Day 8 (Fig 6B; Pearson’s *r=*0.48*, p*=0.004). Thus, the magnitude of the DA reduction induced on the first day of training predicted the probability and speed of escape performance on subsequent days.

**Figure 6.**
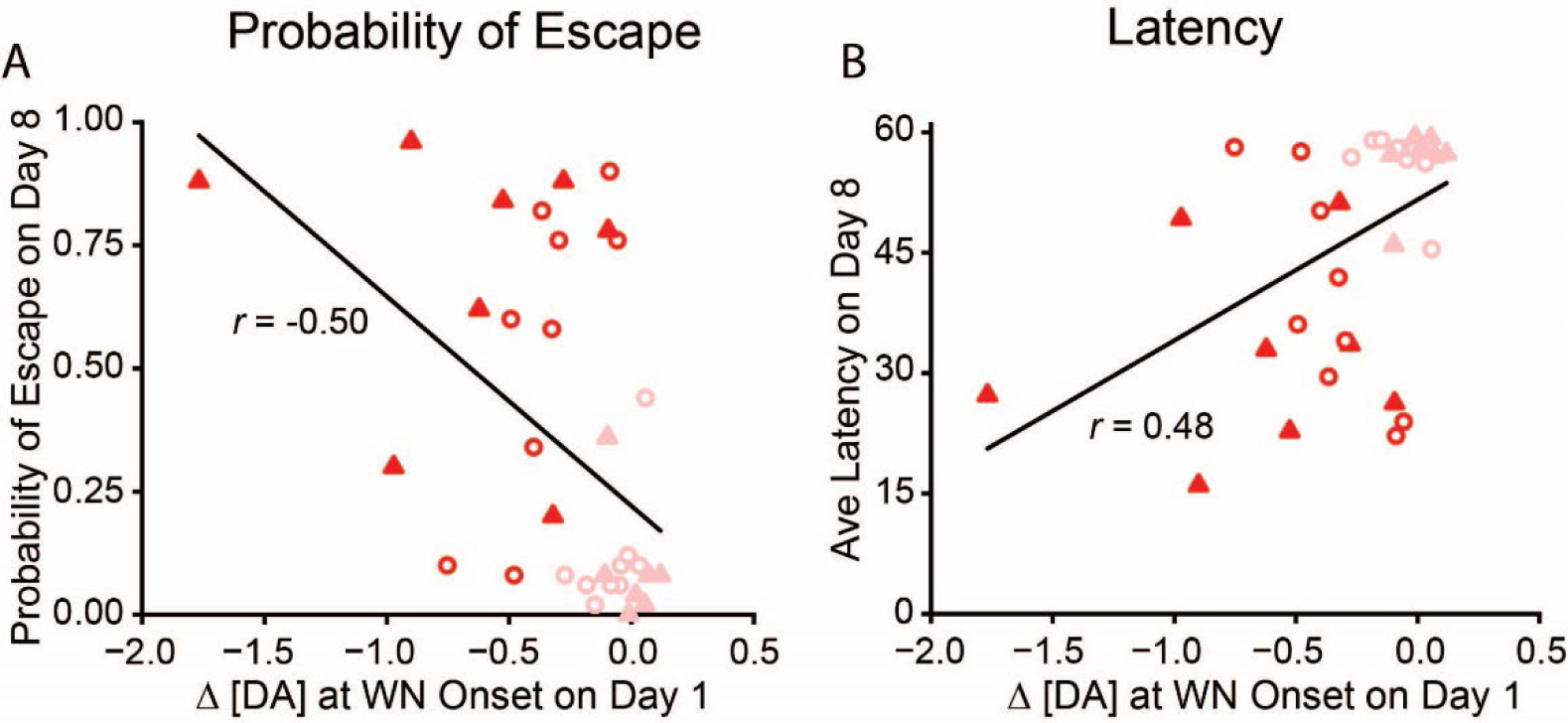
Noise-induced Reductions in Dopamine were Correlated with Increased Escape Behavior. The magnitude of the noise-induced DA change on Day 1 of Negative Reinforcement training had a significant negative relationship with the probability of escape responses on Day 8 Pearson’s r=-0.50, p=0.003. (B) The magnitude of the noise-induced DA change on Day 1 of training had a significant positive relationship with the average escape latency on Day 8 Pearson’s r=0.48, p=0.004. Intense (red); Mild (pink); Male (triangle); Female (circle).

## Discussion

In these studies, we observed that an aversive stimulus reduced NAcC DA and increased motivated behavior in two behavioral designs. Exposure to an aversive WN, but not a mild WN, enhanced cocaine self-administration. This finding adds to a robust literature demonstrating that aversive stimuli and stressors increase the motivation to take drugs in humans^1,3,26,27^ and in rodent models^10,11,28–31^. To examine the generality of this physiological and behavioral phenomenon, we tested the ability of this same intense WN to maintain negatively reinforced behavior. Consistent with the former experiment, intense WN both reduced NAcC DA signaling and promoted motivated escape behavior. It is notable that the experiments presented in this report were powered to detect sex differences, and few behavioral or physiological differences were found. Intense WN consistently decreased DA while promoting cocaine intake and escape behavior in both female and male rats. Based on this, it is reasonable to conclude that this mechanism of aversively motivated behavior is engaged by both sexes to guide behavior.

The observation that an aversive stimulus simultaneously inhibits NAcC DA and stimulates drug taking provides an ethological model for a well-established pharmacobehavioral phenomenon. It has long been known that dopamine receptor antagonists can increase psychostimulant self-administration^32,33^. For example, a moderate dose of the DA receptor antagonist pimozide increases both cocaine and amphetamine self-administration. Following administration of a higher dose, operant responding increases temporarily then diminishes, a phenomenon termed extinction mimicry^17–19^. The former phenomenon resembles what would be observed if the per-unit dose of amphetamine was reduced, and the latter is what would be observed if amphetamine was replaced with saline. The NAcC is a critical site of action for these effects, as pharmacological manipulation of DA receptors in this area controls satiety in cocaine self-administration^20,21^. DREADD inhibition of NAcC-projecting DA neurons produces a similar enhancement of low effort drug taking, showing that the effect is not limited to manipulations of DA receptors^34^. Importantly, the magnitude of post-response DA release in the NAcC has been causally linked to the rate of drug taking. Escalation of cocaine self-administration is accompanied by a muted post-response DA release, and DA terminal stimulation during this period reduces drug intake^35,36^. Taken together, these findings suggest that under pharmacological challenge, animals will work to restore reduced DA signaling. The present report advances this line of investigation by demonstrating that this physiological mechanism could be naturally engaged by a noxious environmental stimulus that reduces DA signaling and promotes both increased drug intake and motivated escape.

Recent studies that have examined NAcC dopamine in aversive learning situations have interpreted their sometimes-conflicting findings as a reflection of various combinations of stimulus salience, hedonic or motivational value, and associative strength^13,37–39^. In the present study, we measured DA during both positive and negative reinforcement and found that the directionality of DA signaling appeared to be dictated by subjective hedonic valence. During positive reinforcement, DA signaling was consistent with its proposed roles in both prediction-error and motivation^40^. When reward delivery was uncertain, sucrose pellet presentation caused an increase in DA, whereas reward omission reduced DA. This pattern is what would be predicted by prediction-error and incentive salience interpretations, provided the reasonable assumption that a variable schedule of reinforcement produces perpetual uncertainty^41^. When switched to a negative reinforcement schedule, the presentation of aversive noise reduced DA, and the termination of that noise produced a timelocked DA release, similar to that caused by the delivery of sucrose. Overall, DA was suppressed by the presentation of an aversive stimulus or the omission of expected reward, whereas DA was released after reward presentation, or the removal of an aversive stimulus. The salience of the event was also meaningful, as the DA decrease induced by an intense noise was greater than that induced by a weaker noise. However, the direction of the DA response was associated with subjective valence rather than salience. Discrepancies between prior studies could be due to variations in animal models or behavioral tasks, but the precise location of observations is also critical^12^. Even within the NAcC, DA neuron terminal activity during aversive learning can vary based on location in functionally significant ways^38^.

We found that the impact of a noise on DA signaling is correlated to its ability to induce escape behavior. This finding is interesting when considered relative to similar studies of cued avoidance learning. Although cues that signal unavoidable aversive outcomes decrease DA in the NAcC^14,37–39^, cues that reliably signal the opportunity to avoid aversive outcomes often increase DA^14,37^, and this positive DA signal can directly modulate avoidance behavior^42^. In the present study, there was no opportunity to completely avoid the aversive outcome, and there was no neutral predictive cue. The escape behavior was triggered by the aversive stimulus, which depressed DA signaling throughout training, even after the acquisition of learning. This suggests a potentially important distinction between escape and avoidance, and is consistent with observations that DA inhibition is associated with, and can induce, escape behavior^14,43^.

In this report, we observed that greater noise-induced reductions in NAcC DA were correlated with increased escape behavior, suggesting that the reduction in DA is a critical component of the motivation to escape. The mechanism through which aversion-induced reductions in DA may support motivated behavior remains unclear. It is possible that reduced DA may differentially affect striatal outputs based on receptor expression. This could include reductions in DA preferentially disinhibiting drd-2 receptor expressing medium spiny neurons (MSNs) in the NAc^44–47^. However, recent reports suggest greater complexity in this system, and that NAc MSN subtypes may instead encode different types of information about an aversive stimulus, such as salience or prediction error^37,48^. Others have also shown that drd-1 and drd-2 receptor activity in the dorsal striatum are both important for the acquisition of negatively reinforced behavior, suggesting that the downstream pathways involve more than just reductions in NAcC DA^49^. The causal role of aversion-induced reductions in DA, as well as the role of the downstream striatal structures that mediate the motivational properties of reduced DA, will be the focus of future studies that could provide new therapeutic targets for treating disorders that are exacerbated by aversive stimuli.

## Author Contributions

EMG, DSW, JRM, MCH, and RAW designed the experiments. EMG, BEC, EG, LV, and DSW performed the surgeries, conducted the experiments, and analyzed the data. EMG, DSW, and RAW wrote the manuscript. JRM, MCH, LV, NP and DSW provided critical feedback on the manuscript.

## Acknowledgments

This work was supported by the Charles E. Kubly Mental Health Research Center and the National Institutes of Health (DA048280 to RAW, MCH, and JRM). The authors would like to thank Talia Lerner, Vaibhav Konanur, Matthew Gelin, Peter Lamberton, Kara Zimolzak and Mitch Roitman for technical assistance.

