## Supplemental Materials for "Aversion-induced drug taking and escape behavior involve similar nucleus accumbens core dopamine signaling signatures"

### Detailed Methodology

**Subjects.** Adult female and male Sprague-Dawley and Long Evans rats (200-350g; Envigo) were used in all experiments. Animals were individually housed on a reverse 12:12 light-dark cycle in a temperature- and humidity-controlled, Association for Assessment and Accreditation of Laboratory Animal Care accredited vivarium. All procedures were approved by the Marquette University Institutional Animal Care and Use Committee. During behavioral training and testing, and for 6 days prior, animals were fed standard chow (LabDiet) once daily to maintain 90% body weight. Water was available *ad libitum* for the duration of all experiments.

**Surgical procedures.** All surgical procedures were conducted under isoflurane (2.0 - 2.5%) anesthesia. For the cocaine self-administration experiment, intrajugular catheters were implanted as previously described[1]. To prepare for photometry recordings, animals were head-fixed for stereotaxic implantation of an optic fiber unilaterally targeting the NAcC and viral injection of AAV5-hSyn-dLight 1.2 or 1.3b (AP: +1.3 mm; ML: +1.3 mm; DV: -6.9 mm; 1µL/10 min; titer = 2.3 or 2.5 x $10^{13}$ molecules/mL). Next, an optic fiber (9 mm length, 400 µm core/430 µm outer diameter with 0.48 numerical aperture, flat tip; Doric) was implanted at the same site. Rats were treated with the anti-inflammatory drug, meloxicam (1mg/kg, s.c.) the day of surgery and for 2 d following surgery to reduce inflammation and postoperative pain. To maintain patency, the catheters were flushed daily with heparinized saline and the antibiotic cephazolin. Rats were allowed to recover for 2-4 weeks before food restriction and behavior.

**Behavioral apparatus.** Subjects were tested in standard operant chambers (Med Associates), interfaced with a computer, and housed in sound attenuating cabinets. Each operant chamber had a dedicated syringe pump for intravenous infusions. Two retractable levers entered the chamber on the right-side wall with a cue light directly above each. A food-pellet dispenser delivered 45-mg sucrose pellets (Bio-Serv) to a recessed foodcup positioned between the two levers. A speaker capable of producing a WN (55 - 90 dB) was located on the left sidewall. The background noise was ~48 dB.

**Photometry apparatus.** Simultaneous recording of dLight fluorescence and background was accomplished using two separate wavelengths of light (465 nm and 405 nm, respectively) provided by an RZ10X processor (Tucker Davis Tech), or by two single wavelength LEDs (Doric) controlled by an external dual channel driver (Doric), which itself was driven by an RZ5P processor (Tucker Davis Tech). Both wavelengths were routed through a dichroic mirror (4-port fluorescence mini cube, Doric) and combined into a single 2-meter jacketed patch cord (400 µm core, 0.48 numerical aperture; Doric). This fiber was secured to the optic fiber implanted in the animal using a ceramic sleeve (Precision Fiber Products) and custom-made screw clamp. This fiber carried both the excitation and emission fluorescence, which were separated by a dichroic mirror that delivered the dLight fluorescence to the RZ10X processor or to a Newport Visible Femtowatt photoreceiver (Doric; delivered by 600 µm core/630 µm outer diameter, 0.48 numerical aperture patch cord, Doric). Recordings occurred using commercially available software (Synapse; Tucker Davis Tech) at 1017.2 Hz. Signals were recorded for at least 5 minutes prior to the beginning of each behavioral session to permit early signal decay.

**Analysis.** Data were extracted using modified Matlab scripts (Tucker-Davis). Both the 405 and 465 nm signals were downsampled by a factor of 100 from the original sampling rate. The processed isosbestic signal was fitted to the excitation signal using a linear fit to correct for signal decay. The dLight excitation signal was then normalized by subtracting the fitted isosbestic from it and dividing the difference by the fitted isosbestic, yielding the ΔF/F.

The ΔF/F signal was normalized for each session for each subject using a pre-program mean Z-score ($Z = (X - \bar{X})/(SD)$, where $\bar{X}$ = mean ΔF/F from a baseline period preceding the onset of behavioral testing, and SD = the standard deviation of the mean ΔF/F from the same period). The response was visualized by aligning the ΔF/F-Z to variables of interest for each behavioral measure using NeuroExplorer 4.0 (Nex Technologies).

**Cocaine-only self-administration**. Eight female and 7 male rats underwent catheter surgery. After recovery and food restriction, rats were trained to press one of two levers for a sucrose pellet reward on an FR1 schedule in computer interfaced operant conditioning chambers enclosed in sound attenuating cubicles (Med Associates). After the acquisition of lever pressing behavior (6-7 1-hr daily sessions), rats were trained to self-administer cocaine on an FR1 schedule. The beginning of each cocaine self-administration session was signaled by the entry of both levers into the box and the illumination of two cue lights. Responses on the active lever resulted in a 2.6-s cocaine infusion (0.5 mg/kg/0.1 ml) accompanied by cue-light offset, active lever retraction, and a 20-s timeout period. The conclusion of the time out period was signaled by cue light onset and extension of the active lever. Responses on the inactive lever resulted in no programed consequences.

**WN testing during cocaine-only self-administration**. After acquisition of stable cocaine self-administration (12-14 daily 2-hr sessions), rats were tested with mild (55 dB) and intense (90 dB) WN. Each WN test session began with a 25 min self-administration period with no auditory stimulation. This was followed by 3 WN test cycles. Each test cycle consisted of two 5-min WN presentations (WN “on” periods) separated by three 5-min quiet periods (WN “off” periods, for a total of 25 min (off, on, off, on, off; see Figure 1). Each cycle was separated by a 1-min break during which both cue lights were extinguished, and both levers were retracted. Each rat experienced 2 intense test days and 2 mild test days (order counterbalanced) for a total of 12 WN exposures of each intensity. These test days were separated by 5 quiet test days featuring the same structure, but no WN presentations. The effect of WN on cocaine self-administration was assessed by subtracting the average infusions during the “on” periods from the average infusions during the “off” periods. On quiet days, time-matched “on” and “off” periods were used for comparison.

**Concurrent food and cocaine self-administration.** A separate cohort of rats (7 male and 5 female Long-Evans) received IV catheter surgeries and were trained to self-administer cocaine as described above with the following exceptions. The animals were initially trained to press two levers for sucrose pellets. After acquisition of lever pressing, all rats received 5 cocaine (0.8 mg/Kg/0.2 ml infusion) self-administration sessions in which only one lever was present. Responses on this lever delivered only cocaine infusions for the remainder of the experiment. For the next 9-10 sessions, the second lever was reintroduced, and rats received simultaneous access to cocaine and sucrose pellets. Both reinforcers were presented on the same schedule with the same response consequences and timeout. The responses were never mutually exclusive. The subjects then received two WN test sessions and two quiet test sessions. Each WN test day consisted of six 10-min intense WN on periods interspersed with 10-min WN off periods. Unlike previous tests, the test sessions began with a WN on period.

**Photometric recording during cocaine-only self-administration**. A separate cohort of rats (6 male and 5 female) received IV catheter and dLight photometry surgeries and were trained to self-administer cocaine without concurrent food as described above. On the penultimate day of cocaine training, the rats were tethered and habituated to the recording situation. If the tether noticeably decreased self-administration, rats were given an additional habituation day. Photometry recordings occurred on 4 nonsequential days featuring 2 quiet test days and 2 intense WN test days (counterbalanced). Test day procedure was identical to the cocaine-only test procedure described above, except the break period between cycles was used to untangle the tether if necessary. Between test days, rats received regular self-administration sessions without the recording tether in order to reduce fatigue. DA changes in response to WN were assessed by comparing the average Z-scored ΔF/F during the 60-s after noise onset (or matched “on” period during quiet days) to a baseline measure taken in the 60-s period immediately before noise onset. DA changes during cocaine infusions were assessed by comparing average Z-scored ΔF/F during the 5-s period immediately after the lever press to a baseline measure taken in the 2-s period starting 3-s before the lever press. To examine the relationship between DA and behavior, a difference score was calculated for each trial by subtracting the average DA during the 60-s baseline period from the average DA during the 60-s post onset period. These difference scores were averaged into 6 two-trial blocks, and compared to the corresponding lever-press difference scores.

**Positive reinforcement.** To examine DA signaling during positive and negative reinforcement, 23 female and 21 male rats received dLight photometry surgery. After recovery and food restriction, rats began lever training in which each of two levers were presented independently and reinforced with a sucrose pellet on an FR1 schedule until the subjects received 50 reinforcers. Following 4 daily sessions of lever training, both levers were extended simultaneously and rewarded on independent VI30-s schedules in a 20-min session. This schedule continued for 3 days before progressing to a VI90-s schedule for 6 daily 45-min sessions. Photometry recordings were taken on an early and a late day of VI90 positive reinforcement training. Changes in DA were assessed by subtracting the average fluorescence in the 5-s period immediately following the lever press from the 2-s period immediately preceding the lever press. Timestamps with corresponding 405 signal or ΔF/F-Z scores that exceeded 4 standard deviations from the average 405 signal or ΔF/F-Z score were excluded from analysis.

**Negative reinforcement.** After the final day of VI90 food training, a subset of the same rats (22 female, 19 male) received negative reinforcement lever training. In this design, operant responding on the left or right lever (counterbalanced) was reinforced by the termination of an aversive WN. Rats went through the negative reinforcement acquisition phase in which they were presented with 60 trials of WN each day. Rats were either trained with an intense (90 dB) or a mild (55 dB) WN. Each trial began with WN onset. After 5-s, both levers extended but responses on either lever had no consequence. Beginning 10-s after the onset of the WN, a response on the active lever resulted in the termination of the WN for 20-s. Then the next trial began with the onset of the WN. Choosing the inactive lever resulted in retraction of both levers for 5-s, but the WN persisted. If the rat failed to terminate the WN, it would persist for a total of 60-s, followed by a 6-s timeout period before the start of the next trial. This training continued for 5 days. Then, 10 pseudorandomly interspersed quiet trials were introduced to the NR session such that the rat experienced 50 trials of WN and 10 trials in which no noise was presented. Quiet trials were always 60-s, and choices on either lever only resulted in lever retraction for 5-s. Rats received three days of training with the quiet trials included. Probability of response for a given lever was determined by dividing the number of responses by the number of presentations of the WN. Latency was defined as the average time to terminate WN across the session. Photometry recordings were taken on an early and a late day of initial NR training as well as during the middle day of the training with quiet trials. Some animals were excluded from analyses because they did not yield data on one or more days of testing (Table S1). Changes in DA in response to the operant response were assessed by subtracting the average fluorescence in the 5-s period immediately following the lever press from the 2-s period immediately preceding the lever press. Changes in DA in response to WN were assessed by subtracting the average fluorescence in the 5-s period immediately following the WN onset from the 5-s period immediately preceding WN onset. Timestamps with corresponding ΔF/F-Z scores that exceeded 4 standard deviations from the average ΔF/F-Z score were excluded from analysis.

**Histology.** Following the final day of behavioral testing, animals with fiber photometry surgeries were perfused with saline and 4% paraformaldehyde. Brains were extracted and stored overnight in 4% paraformaldehyde then transferred to a cryoprotectant 30% sucrose solution. After 3-4 days in this solution, brains were flash frozen with 100% ethanol and dry ice. Brains were sliced at 35uM on a cryostat, then were processed to assess dLight fluorescence. After mounting each slice on a slide, brains were imaged at 10x on a fluorescent microscope (Keyence) to verify viral expression and fiber placement (Fig S1A-B).

**Expanded Results**

**Histological Verification of Fiber Photometry Recordings**

For all experiments, following the final day of behavioral testing, animals with fiber photometry surgeries were processed to verify viral expression and fiber placement.

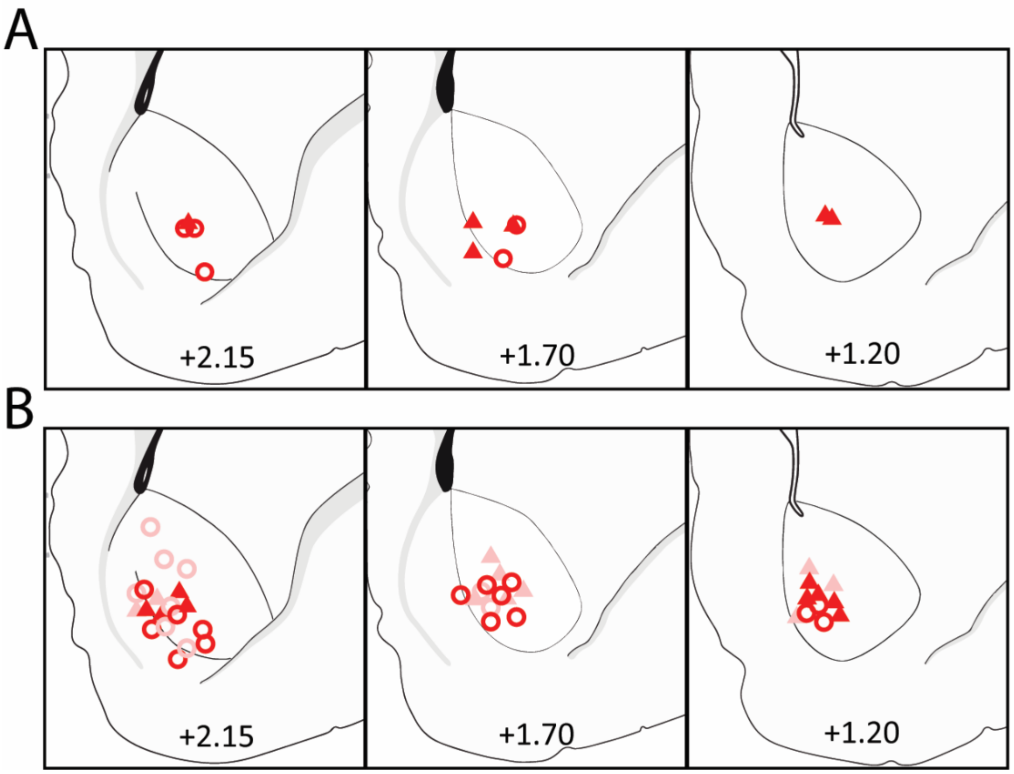


Figure S1. Histological Verification of NAc DA recordings. (A) Optic fiber placements for the cocaine photometry experiment. (B) Optic fiber placements for the negative reinforcement study. Placements were verified to be in the NAc core or border between the NAc core and shell for all photometry experiments. Female placements are depicted as circles. Male placements are triangles. The intense white noise condition is depicted in red; Mild is pink.

**Operant Response Data for Cocaine Self-Administration**

Responding on the inactive, cocaine, and food paired levers was recorded during testing.

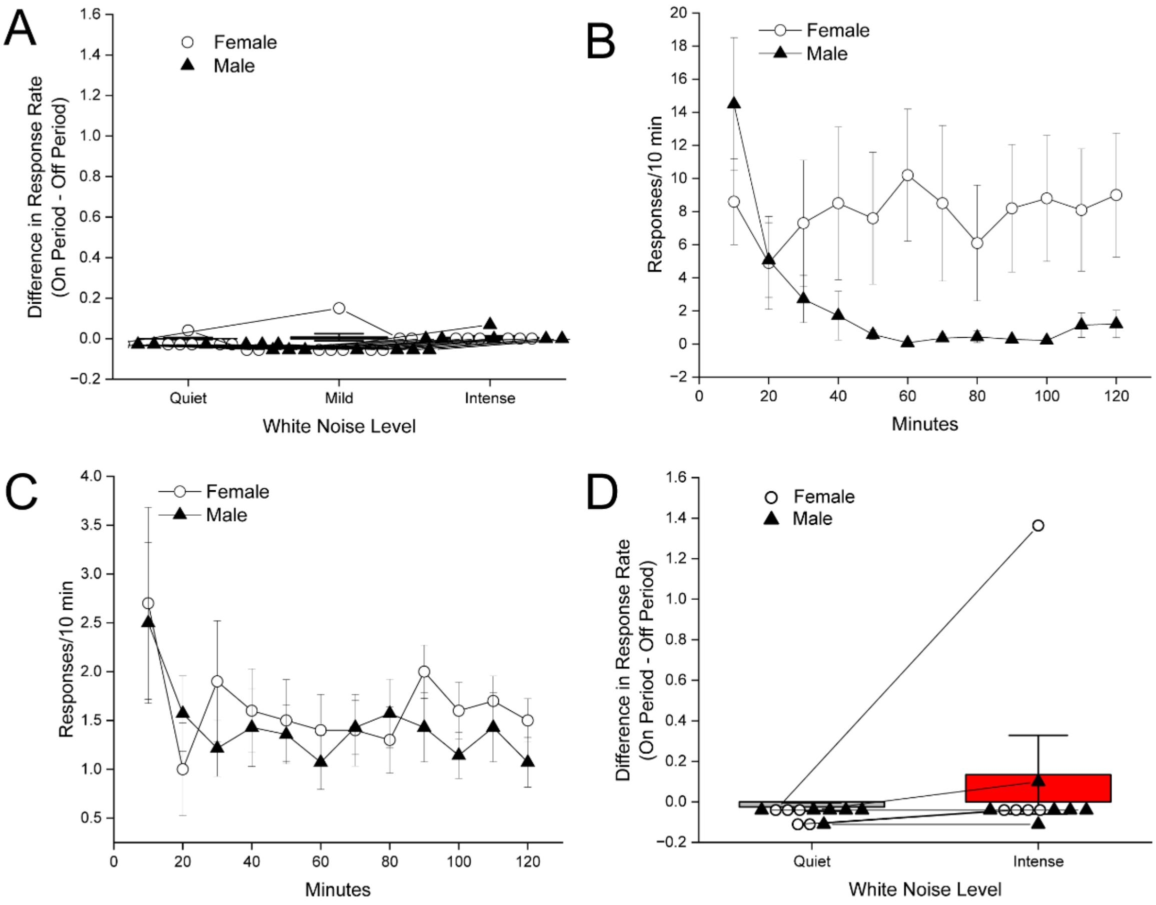


Figure S2. Responding on Inactive, Cocaine-, and Food-Paired Levers. Inactive lever responses were analyzed for the cocaine self-administration studies. (A) Cocaine-only self-administration: Mean difference in response rate on the inactive lever during quiet, intense (90 dB), and mild (55 dB) noise on periods vs off periods. +/- SEM (B) Concurrent food and drug self-administration: Mean food response rate across quiet test sessions during. +/- SEM (C) Concurrent food and drug self-administration: Mean cocaine response rate across quiet test sessions. +/- SEM (D) Cocaine-only self-administration photometric recordings: Mean difference in response rate on the inactive lever during quiet or intense (90 dB) noise on periods vs off periods +/- SEM.

**Expanded Negative Reinforcement Data**

<u>After Several Days of Exposure, Mild WN Reduced NAc Dopamine</u>

Rats underwent 8 days of negative reinforcement testing, including three days with fiber photometry recordings of DA. On Day 4 (Fig S3A-B), both intensities of white noise produced a reduction in DA, with an interaction between intensity and epoch (ANOVA: $F(1,33) = 11.31$, $p=0.002$). Intense white noise continued to produce a large reduction (PLC: $F(1,33) = 77.23$, $p<0.001$). By this time, the mild white noise produced a smaller, but significant, reduction in dopamine (PLC= $F(1,33) = 10.37$, $p=0.003$). Additionally, this analysis revealed a significant main effect of Sex ($p=0.025$), showing that females had higher levels of DA overall, regardless of intensity or time relative to WN onset.

Comparable analyses were conducted for Day 7 (Fig S3C-D). Both intensities of white noise continued to reduce dopamine, though there was an interaction between intensity and epoch ($F(1,31) = 6.07$, $p=0.002$). Intense white noise continued to produce a large reduction in dopamine ($F(1,31) = 92.93$, $p<0.001$). The mild white noise, similar to Day 4, continued to produce a reduction in dopamine as well ($F(1,31) = 30.67$, $p <0.001$). This analysis also revealed a Sex x Intensity interaction ($F(1,31) = 11.02$, $p=0.003$) reflecting reduced DA in the female rats assigned to the intense noise condition.

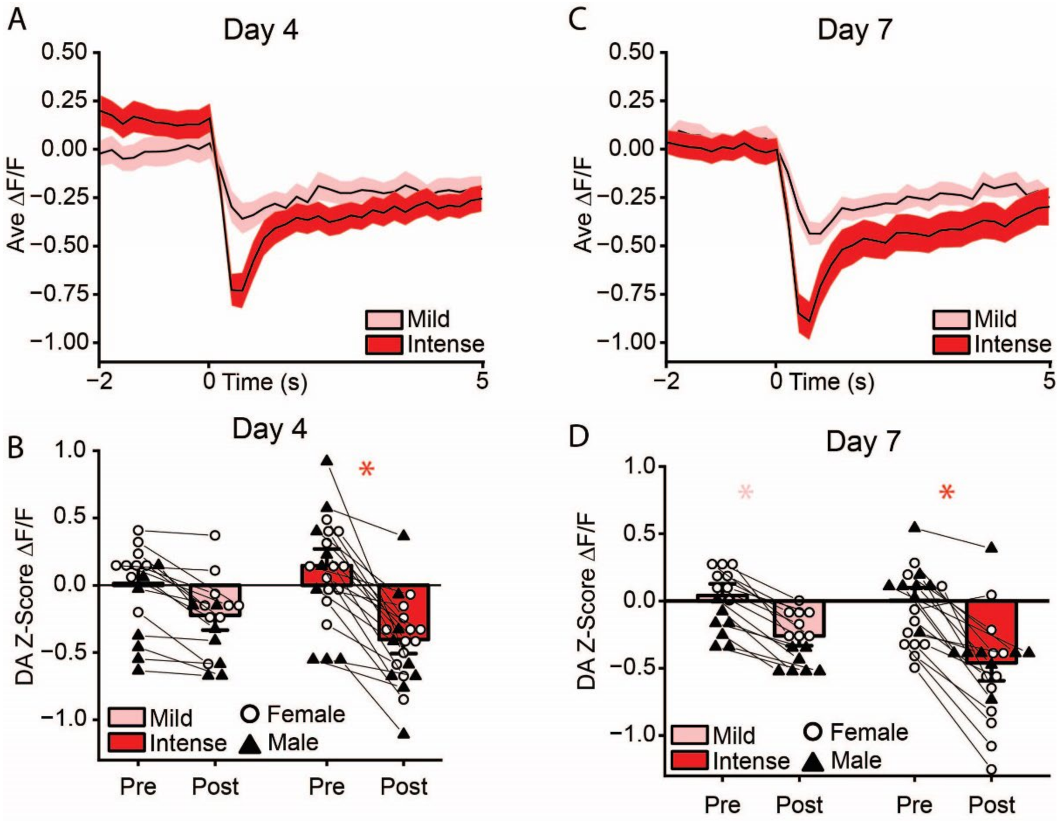


Figure S3. After Days of Exposure, Mild WN Reduced NAc Dopamine. (A,C) Mean +/-SEM Dopamine concentration timelocked to onset of Intense (red) or Mild (pink) WN on the fourth (A) and seventh (C) day of negative reinforcement training. (B,D) Dopamine concentration in the 2s prior to (pre) vs 5s after (post) WN onset. Day 4, Intense: $F(1,31) = 77.23$, $p<0.001$. Day 4, Mild: $F(1,33) = 10.37$, $p<0.01$. Day 7, Intense: $F(1,31) = 92.93$, $p<0.001$. Day 7 Mild: $F(1,31) = 30.67$, $p<0.001$)

<u>Following Chronic Exposure and Retraining, Mild WN Maintained Negatively Reinforced Behavior</u>

Our overarching hypothesis is that aversion-induced reductions in DA may promote behavior through motivational or learning mechanisms. Consistent with this, mild WN failed to reduce DA early in training and correspondingly failed to support the acquisition of escape behavior. However, following repeated exposure, mild WN did significantly reduce DA. This created an opportunity to determine if this decrease in NAc dopamine reflected a sensitization to the aversive properties of the mild WN such that the same stimulus could now support the acquisition of negatively reinforced behavior. To test this, a subset of the rats (n=8) that received mild WN received an additional re-training period consisting of three days of lever responding for sucrose reinforced on a VI90 schedule followed by 8 days of NR training with mild WN, all of which were structured identically to the initial training sessions (Fig S4A). Due to the smaller sample size, the retraining portion of the results was not powered to detect sex-differences, though an equal number of male and female rats contributed behavioral and physiological data to this analysis.

As during initial training, rats reduced responding on the inactive lever during negative reinforcement retraining (Fig S4B), verified by a t-test of their responding on the last day of food re-training and the last day of NR re-training ($t(8)= 2.52$, $p=0.040$). In contrast, following mild WN exposure and retraining, the presentation of mild WN caused rats to maintain responding on the active lever throughout negative reinforcement (Fig S4B; $t(7)= 0.238$, $p=0.818$). This change in escape behavior was verified by within-subject comparisons of the last day of initial training (NR8) and the last day of retraining (NR16) (Fig S4C). A t-test was conducted on these days which and revealed an increase in active responses on NR16 relative to NR8 ($t(7)= -3.91$, $p=0.006$), but no change in inactive responses, $t(7)= -1.58$, $p=0.158$). To further test the within-subjects change following noise exposure and retraining, a t-test was conducted on the average latency to escape the WN on the last day of initial training vs the last day of retraining (Fig S4D). This analysis showed that rats terminated the WN more quickly on the last day of retraining (NR16) than on the last day of initial training (NR8; $t(7)= 3.48$, $p=0.010$).

To verify that the mild WN continued to reduce DA after retraining, we compared DA measures on the first day of initial training with those on the first day of the retraining (NR1 vs NR9). Rats showed a reduction in dopamine on the retraining day, but not on the initial training day (Fig S4E-F; ANOVA: $F(1,14) = 6.36$, $p=0.024$). On the first exposure to mild white noise, rats did not show a reduction in dopamine at the onset of the noise (PLC: $F(1,14) = 0.51$, $p=0.485$). However, after substantial exposure and re-training on lever pressing, rats showed a significant reduction in dopamine (PLC: $F(1,14) = 47.23$, $p<0.001$).

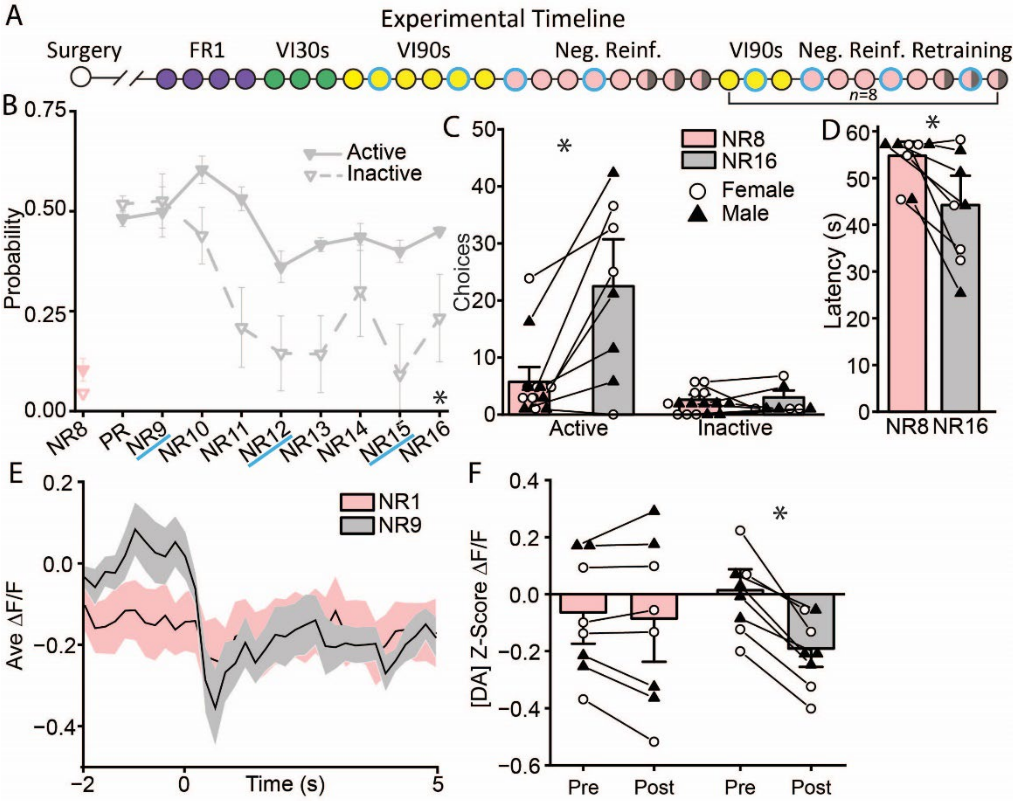


Figure S4. Following Chronic Exposure and Retraining, Mild WN Sustained Negatively Reinforced Behavior. (A) Experimental Timeline. A subset (n=8) of rats that received mild WN in the previous experiment were re-trained on positive reinforcement, then transitioned back to negative reinforcement. (B) The probability of a response on the active (solid) or inactive (open) levers across days of retraining. Consistent with initial training, relative to the probability of choice on the last day of re-trained positive reinforcement, rats pressed less on the inactive lever by the end of retraining ($t(7) = 2.52$, $p=0.040$). In contrast, following mild WN exposure and retraining, the presentation of mild WN caused rats to maintain their response on the active lever relative to the re-trained positive reinforcement, ($t(7) = 0.238$, $p=0.818$). (C) Within-subject comparison of active and inactive choices on the last day of initial training (NR8, pink) vs the last day of retraining (NR16, gray). Active: $t(7) = -3.91$, $p=0.005$), Inactive: $t(7) = -1.58$, $p=0.157$. (D) Within-subject comparison of latency on the last day of initial training (NR8, pink) vs the last day of retraining (NR16, gray): $t(7) = 3.48$, $p=0.010$). (E) Mean +/-SEM Dopamine concentration timelocked to onset of Mild WN on the first day of initial training (NR1, pink) and the last day of retraining (NR9, gray). (F) Dopamine concentration in the 2s prior to (pre) vs 5s after (post) WN onset. NR1: $F(1,14) = 0.51$, $p=0.485$. NR9: $F(1,14) = 47.23$, $p<0.001$.

The unexpected sensitization to the mild WN further illustrates the relationship between DA inhibition and escape behavior. Initially, mild WN did not inhibit DA or support negative reinforcement learning. However, over the course of several days of exposure, rats began to show a moderate reduction in DA in response to the mild WN, and upon retraining the stimulus did support negative reinforcement learning. This apparent sensitization effect was surprising, since mild WN is routinely used during behavioral testing as ambient noise, although in most studies WN is not presented intermittently as in this report[3–7]. However, the development of increased sensitivity to the aversive properties of electric shock following repeated exposure is well established[8,9]. Most relevant to this report, this effect is not exclusive to intensely aversive stimuli, as an increased behavioral response is also observed following repeated exposure to a mild 0.25-mA shock[10].

**Table S1: Group sizes for all analyses.**

| Figure | Analysis | Intense | | Mild | |
|---|---|---|---|---|---|
| | | Female | Male | Female | Male |
| 1E | Cocaine infusion rate under various noise conditions | 8 | 7 | | |
| 2A | Effect of noise on cocaine vs food | 5 | 7 | | |
| 2B | Cocaine infusion rate under quiet vs. intense noise conditions during photometry | 5 | 6 | | |
| 2E | Change in dopamine around WN onset | 5 | 6 | | |
| 2F | Change in dopamine around WN onset | 5 | 6 | | |
| 2G | Cocaine intake correlations | 4 | 6 | | |
| 3C | Change in dopamine around lever press for cocaine | 5 | 6 | | |
| 3D | Change in dopamine around lever press for cocaine | 5 | 6 | | |
| 4B | Lever responses over days of NR | 11 | 7 | 8 | 9 |
| 4C | Average NR latency | 13 | 10 | 9 | 9 |
| 4D-E | Dopamine responses to positively reinforced behavior | 14 | 9 | 9 | 8 |
| 4F-G | Dopamine responses to negatively reinforced behavior | 10 | 9 | 9 | 7 |
| 5C-D | Naïve dopamine responses to onset of white noise | 10 | 9 | 9 | 7 |
| 6A-B | NR-DA correlations (Day 3) | 10 | 9 | 9 | 7 |
| 6C-D | NR-DA correlations (Day 8) | 9 | 9 | 9 | 7 |
| S3A-B | Dopamine responses to onset of white noise after moderate exposure | 12 | 9 | 9 | 7 |
| S3C-D | Dopamine responses to onset of white noise after substantial exposure | 11 | 8 | 9 | 7 |
| S4 | Retraining of mild negative reinforcement, dopamine and behavior | | | 4 | 4 |

Table S1 Legend.
Group sizes that contributed to specific analyses are presented. Animals did not contribute data to specific analyses if there was a technical issue with recording on the test day (e.g. inadequate signal quality), experimenter error, or behavioral performance was unexpectedly inhibited (<10% of typical responding).
